# The Genomic Architecture of Local Adaptation in Two Connected Populations of Three-Spined Stickleback

**DOI:** 10.1101/2025.10.21.683682

**Authors:** Sann Delaive, Nicolas Derôme, Sam Yeaman

## Abstract

Populations often adapt to their local environments despite the homogenizing effects of gene flow, but the genomic mechanisms enabling this process remain unclear. Theory predicts that adaptive divergence under high connectivity is favored when beneficial alleles cluster in regions of reduced recombination, a pattern that can be reinforced by structural variants (SVs). We investigated this in three-spined sticklebacks (Gasterosteus aculeatus) from the St. Lawrence Estuary, where distinct freshwater and marine ecotypes meet and interbreed along a short ecological gradient. Using long- and short-read whole-genome sequencing, we mapped fine-scale recombination landscapes, catalogued SVs, and examined their relationship with adaptive genomic regions. Recombination landscapes differed between populations, with population-specific shifts in recombination rate estimated by an LD-based method. Putatively adaptive regions were not confined to low-recombination regions, yet SVs (inversions, insertions, and deletions) frequently coincided with local recombination suppression and elevated differentiation, suggesting they may contribute to local adaptation. Differentiated regions also overlapped disproportionately with previously-identified regions involved in repeated local adaptation across the species range, which tended to be strongly enriched on chromosomes IV, VII and XXI. These repeated regions were associated with lower recombination rates, suggesting that recombination suppression may contribute to their reuse across populations. As found in stickleback populations from other regions, the St. Lawrence populations exhibit elements suggestive of concentrated architectures clustered in a few genomic regions, along with relatively diffuse patterns of highly differentiated regions distributed genome-wide, across a wide range of recombination rates. These results highlight the intertwined roles of recombination variation and structural variation in shaping evolutionary trajectories in connected populations.

**Article Summary:** This study investigates how recombination and structural variants shape local adaptation in the three-spined stickleback (*Gasterosteus aculeatus*). Using long- and short-read genome sequencing, the authors compared recombination landscapes and structural variants between marine and freshwater populations from the St. Lawrence Estuary. They found population-specific changes in recombination rate and frequent overlap between structural variants, reduced recombination, and genomic regions showing high differentiation. These findings suggest that variation in recombination and structural variants jointly influence how adaptation proceeds in connected populations, providing new insights into the genomic mechanisms that maintain diversity despite ongoing gene flow.

## Introduction

Understanding how populations adapt to their local environments remains a central question in evolutionary biology. Local adaptation occurs when resident genotypes exhibit higher fitness in their native environments compared to non-resident genotypes (Kawecki and Ebert, 2004). This process generates intraspecific diversity, potentially enhancing a species’ evolutionary potential in the face of environmental change (Hu *et al*., 2021).

Although driven by natural selection, local adaptation can be constrained by gene flow, which homogenizes allele frequencies between populations and replace adaptive alleles by neutral or even introduces locally maladaptive alleles (Feder, Egan and Nosil, 2012). Yet, empirical evidence shows that local adaptation can persist even under high gene flow (Limborg *et al*., 2012; Pespeni and Palumbi, 2013; Muir *et al*., 2014). For instance, Trinidadian guppies have maintained locally adapted phenotypes despite repeated introductions of individuals experiencing different predation regimes (Fitzpatrick *et al*., 2015). Cases such as this raise a key question: which mechanisms might enable local adaptation to persist despite ongoing gene flow?

One explanation lies in the genomic architecture of adaptation, which encompasses the number, effect size, location, and linkage of adaptive loci. Under high gene flow, theory predicts that selection should favor clustered architectures in which adaptive alleles are tightly linked and thus inherited together (Yeaman and Whitlock, 2011; Yeaman, 2022). Recombination, which reshuffles alleles and breaks linkage, plays a central role in shaping these architectures. While it promotes genetic diversity, recombination can also disrupt favorable allele combinations. Thus, local adaptation is often facilitated when recombination is locally reduced, preserving clusters of adaptive loci (Tigano and Friesen, 2016; Peñalba and Wolf, 2020). Identifying how and where recombination landscapes vary between populations in different evolutionary scenarios and assessing the role of such variation on the genomic architecture of local adaptation is required to better understand what enables local adaptation to persist in the face of gene flow.

Recombination rates vary widely across genomes and among populations, influenced by chromosomal features (e.g., relatively low in centromeres and higher near telomeric regions), GC content (Fullerton, Bernardo Carvalho and Clark, 2001), epigenetic modifications (Shilo *et al*., 2015), and structural variation. In particular, large structural variants (SVs)—including inversions, insertions, deletions, and duplications—can locally suppress recombination by disrupting homologous pairing and crossover formation (Völker *et al*., 2010; Morgan *et al*., 2017) and by yielding unbalanced karyotypes in recombinants (Kirkpatrick, 2010). As a result, SVs have emerged as strong candidates for facilitating local adaptation under gene flow (Jay, Aubier and Joron, 2018; Wellenreuther *et al*., 2019).

Inversions, detectable through patterns of linkage disequilibrium (LD) or local PCA (Ma and Amos, 2012; Mérot, 2020; Euclide *et al*., 2024), have received particular attention, with clear examples of inversion-mediated local adaptation in plants (Soudi *et al*., 2023), insects (Mérot *et al*., 2021), fishes (Meyer *et al*., 2024), and mammals (Harringmeyer and Hoekstra, 2022). However, the adaptive role of other SV types—such as copy-number variants (CNVs), including insertions, deletions, and duplications— is also increasingly recognized. For example, CNVs associated with thermal adaptation have been reported in Atlantic capelin (Cayuela *et al*., 2020), American lobster (Dorant *et al*., 2020), pika (Sjodin *et al*., 2024), and common ragweed (Wilson *et al*., 2025). Despite their importance, non-inversion SVs remain underexplored due to historical challenges in detection. Advances in long-read sequencing now allow comprehensive characterization of all SV types within a single study (Mérot *et al*., 2020; Lecomte *et al*., 2024), avoiding biases inherent to indirect detection methods (Noor and Bennett, 2009).

The three-spined stickleback (*Gasterosteus aculeatus*) is a well-established model for studying local adaptation. Having repeatedly colonized freshwater habitats from marine ancestors across the Northern Hemisphere (Fang *et al*., 2018), sticklebacks exhibit repeated phenotypic and genomic changes associated with freshwater adaptation. Resources such as a high-quality reference genome (Nath, Shaw and White, 2021), a pedigree-based recombination map (Rastas *et al*., 2016), and extensive ecological and genomic data (Hohenlohe and Magalhaes, 2020) have enabled detailed study of adaptive divergence between marine and freshwater populations. Freshwater adaptation in sticklebacks often displays a concentrated genomic architecture, characterized by a few large-effect loci embedded within “islands of differentiation”, such as the well-known *Eda* locus controlling lateral plate number (Jones *et al*., 2012; Roberts Kingman *et al*., 2021), as well as many smaller-effect loci scattered across the genome. These islands are often reused across independent freshwater colonization events (Hohenlohe *et al*., 2010; Jones *et al*., 2012; Roberts Kingman *et al*., 2021), likely because they contain alleles maintained as standing genetic variation in the ancestral marine population (Nelson and Cresko, 2018). Multiple studies have also demonstrated the role of different structural variants in freshwater adaptation (Chain *et al*., 2014; Rodríguez-Fuentes *et al*., 2025). Moreover, Shanfelter, Archambeault and White (2019) have demonstrated the potential for recombination landscape to vary greatly between a freshwater and a saltwater population. All these characteristics make the three-spined stickleback a suitable species to study the relationship between recombination, structural variation and their impact on the genomic architecture of local adaptation.

In this study, we used long- and short-read whole-genome sequencing data from two geographically close populations of three-spined sticklebacks, occupying different salinity environments in the St. Lawrence Estuary to investigate the genomic architecture of local adaptation in the presence of gene flow (Delaive *et al*., 2025). We first tested whether observed recombination landscapes estimated based on variation in LD differ between freshwater and marine populations and identified where along the genome such differences are most pronounced. We then evaluated whether SVs contribute to these patterns by analyzing their association with population-specific shifts in recombination. Finally, we asked whether recombination landscape divergence and regions of suppressed recombination tend to coincide with elevated genetic differentiation, as expected if recombination and structural variation jointly shape the retention of locally adaptive variants. Together, these analyses provide new insights into how recombination and SVs interact to shape the genomic architecture of local adaptation in the face of gene flow.

## Materials and Methods

### Study System

We analyzed a subset of the whole-genome resequencing dataset previously published in Delaive *et al*. (2025). In brief, genomic DNA was extracted from fin tissue and sequenced on an Illumina NovaSeq 6000 platform (PE150), targeting ∼15× coverage per individual. After quality control, reads were aligned to the stickleback reference genome and SNPs were called using a standardized pipeline that included duplicate removal, local realignment, and filtering on coverage, genotyping success, and minor allele frequency (MAF > 0.05). The resulting dataset comprised 2,332,752 high-quality SNPs genotyped across 387 three-spined sticklebacks sampled along a salinity gradient in the St. Lawrence Estuary, Canada. For the purposes of this study, we excluded individuals from Baie-Saint-Paul, a genetically distinct population that could bias comparisons between the primary marine and freshwater groups. The final dataset thus consisted of 359 individuals: 89 from a fluvial population and 270 from a marine population. The fluvial population included individuals sampled from three sites in the fluvial estuary (low salinity), whereas The marine population encompassed individuals from eight sites distributed across the middle and marine sections of the estuary, where salinity progressively increases to ∼30 ppt (Figure 1).

**FIGURE 1.**
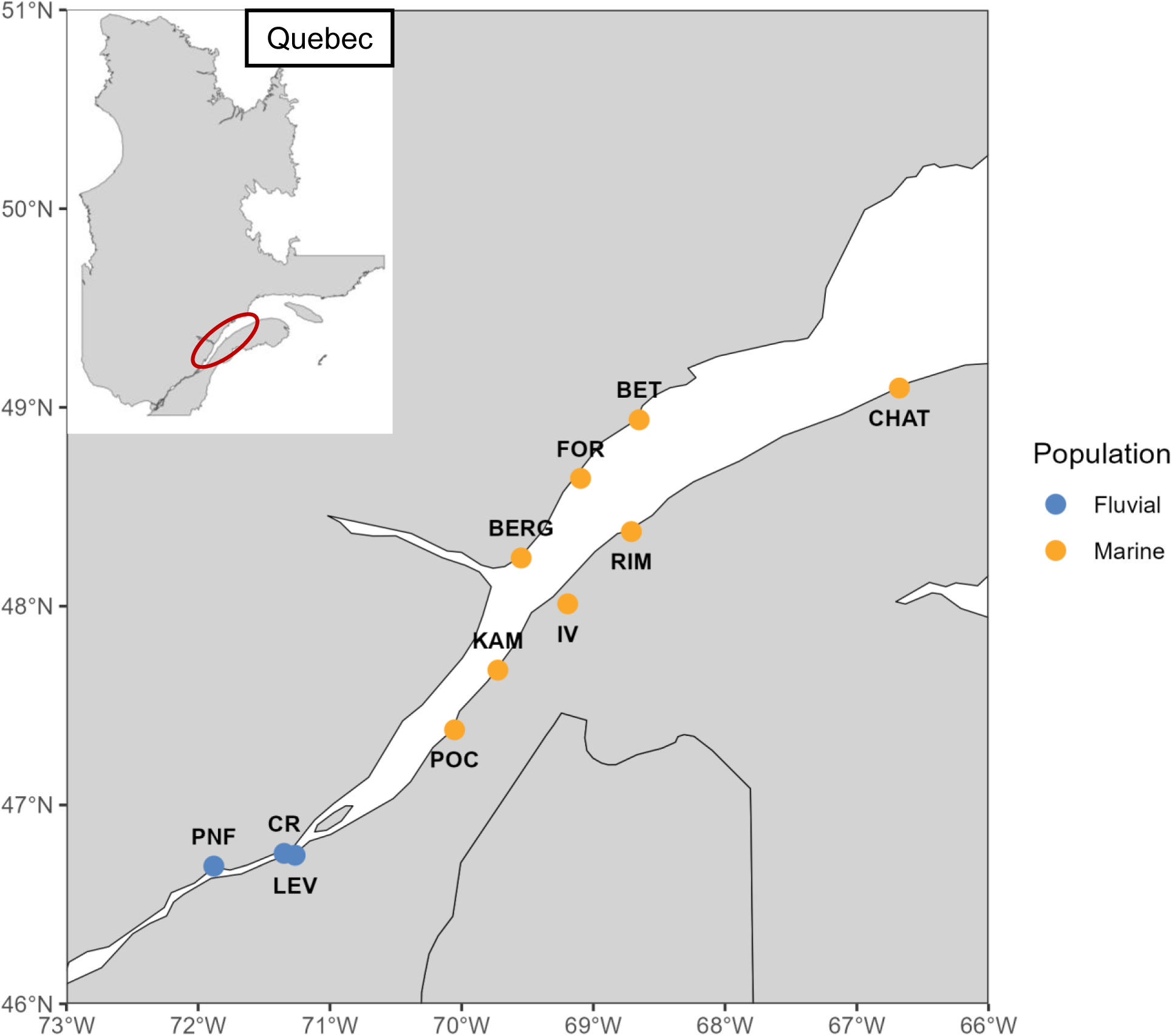
Sampling locations of three-spined stickleback (*Gasterosteus aculeatus*) in the St. Lawrence Estuary. A total of 40 individuals with a balanced sex ratio were collected from 8 sites in the marine and middle estuary, representing the marine population as defined in Delaive et al. (2025), and from 3 sites in the fluvial estuary, corresponding to the fluvial population. Sampling locations are distinguished by colour according to population. Location abbreviations: PNF, Portneuf; CR, Cap-Rouge; LEV, Lévis; POC, La Pocatière; KAM, Kamouraska; IV, Isle-Verte; BERG, Grande-Bergeronnes; RIM, Rimouski; FOR, Forestville; BET, Betsiamites; CHAT, Cap-Chat.

The generation of the dataset is described in greater detail in Delaive *et al*. (2025).

### Construction of Recombination Maps

We reconstructed fine-scale recombination maps using Pyrho (Spence and Song, 2019), which models linkage disequilibrium (LD) using fused-LASSO regression while incorporating historical changes in effective population size (Ne).To provide this information to Pyrho, population-specific Ne trajectories were estimated with SMC++ v1.15.4 (Terhorst, Kamm and Song, 2017), based on randomly subsampled sets of six individuals per population. VCF files were converted using *vcf2smc*, and replicate histories were combined into composite likelihoods to improve inference. Analyses assumed a mutation rate of 4.56 × 10⁻ ⁹ mutations per site per generation (Zhang *et al*., 2023) and a generation time of one year (Reid, Bell and Veeramah, 2021). Because demographic inference relies on patterns of heterozygosity and coalescent events, runs of homozygosity longer than 150 kb were masked prior to inference to avoid overestimation of Ne changes.

To account for within-population heterogeneity, especially in the marine group that encompasses a larger geographical area, we divided each population into two replicate subsets by randomly selecting 42 individuals. We repeated this process three times to create six subsets by population, all comprising a different combination of individuals. The fluvial population was fully included in all replicates, while the marine population subset replicates were drawn from a pool of 84 individuals sampled across six sites to compare similar sample sizes. This resulted in twelve recombination map subsets in total.

Recombination maps were generated using Pyrho following consistent hyperparameter settings across replicates and populations. Lookup tables, representing expected patterns of LD under a specified demographic model and a range of recombination rates, were generated using *--make_table* with a diploid sample size of 42 and a Moran size of 110. The Moran size defines the number of lineages used in the Moran model, a forward-time approximation of the coalescent with recombination and affects the precision of LD expectations under a given demographic history (Kamm *et al*., 2016). Window size and block penalty parameters were optimized using --*hyperparam* by minimizing the L2 norm, defined here as the sum of squared differences between observed and expected LD values. The block penalty determines the smoothness of the recombination landscape by penalizing excessive changes in estimated rates between adjacent genomic windows. Based on this optimization, a block penalty of 15 was applied for all chromosomes, with an optimal window size of 90 SNPs for most chromosomes and 30 SNPs for chromosomes XI and XX. Recombination rates were then estimated with Pyrho’s optimize function using the *--fast-missing* flag. Estimates were obtained separately for each of the six subsets per population, and chromosome-level outputs were merged to produce genome-wide, per-base, per-generation recombination maps for each population.

### Comparisons of recombination maps

To quantify divergence in recombination landscapes between freshwater and marine populations, we applied two complementary metrics introduced by Talbi, Turner and Malinsky (2025): the Population Recombination Divergence Index (PRDI) and a window-based measure of recombination rate dissimilarity, hereafter referred to as Δ*r*. While the PRDI quantifies the broad-scale divergence in recombination landscapes of two populations that cannot be explained by demographic differences alone, the Δr identifies fine-scale patterns of divergence based directly on recombination rate variation.

The PRDI is defined as the difference between two metrics: *me* (empirical divergence) and *ms* (simulated divergence under neutrality). To calculate it, raw recombination rates obtained from pyrho were first averaged in non-overlapping 2 kb windows using the *bedtools map* function. We then assessed the consistency of recombination rate estimates within and between populations by computing pairwise Spearman correlations across subsets using 5 Mb sliding windows along each chromosome. For each window, we calculated all within-population correlations (among freshwater subsets and among marine subsets) and all between-population correlations (across freshwater and marine subsets). The metric *me* was defined as the difference between the lowest median in all within-population correlation and the median between-population correlation.

To establish a neutral expectation (*ms*), we performed coalescent simulations using msprime v1.3.3 (Baumdicker *et al*., 2022). We simulated 84 diploid individuals, partitioned into two groups of 42 to mimic freshwater and marine populations. Simulations assumed a shared recombination landscape across populations, parameterized using the empirically inferred freshwater recombination map (results were unchanged when using the marine map instead). Demographic parameters, including effective population sizes and split times, were informed by SMC++ estimates. For each simulated dataset, recombination maps were reconstructed using the same Pyrho pipeline applied to the empirical data. The *ms* metric was then calculated from the simulated maps following the same procedure as for *me*, and PRDI was defined as the difference between *me* and *ms*.

To further characterize regions where recombination divergence is more pronounced between freshwater and marine populations, we looked for 100 kb windows where the Manhattan distance between two subsets from different populations (between-population Δr) exceeds the Manhattan distance between two subsets from the same population (within-population Δr). Each Δr (within and between) was calculated as the pairwise Manhattan distances in non-overlapping 100 kb windows, each composed of 50 adjacent 2 kb bins. This Manhattan distance was then scaled by the number of bins, and log transformed to stabilize variance. For each 100 kb window, we calculated the median within-population and between-population Δr. Windows where the between-population Δr exceeded three standard deviations above the genome-wide mean of within-population Δr were considered Δr outliers.

Recombination rate varies along the genome and can happen relatively frequently in some regions (hotspots) and rarely in others (coldspots). To understand if these particular regions are conserved between populations, we identified recombination hotspots and coldspots by population using a threshold-based approach. For each window in each empirical recombination map, we computed the local background mean and standard deviation of log-transformed recombination rates within a ±20 kb window. Windows exceeding three standard deviations above or below the local mean were classified as hotspots or coldspots, respectively. Adjacent significant windows were merged with a 1 kb maximum gap using bedtools *merge* to reduce redundancy. Hotspots longer than 5 kb were removed as they are subject to known bias (Hoge *et al*., 2024). We then used bedtools *intersect* to identify hotspots and coldspots consistently shared within and between populations, providing a genome-wide view of recombination variability across the estuary gradient.

### Correlation between linkage map and observed recombination estimates

To mitigate potential circularity in our analyses, where strong selection can locally increase LD around F_ST_ outliers, thereby biasing downward the estimation of observed recombination rates inferred from LD-based methods like Pyrho, we complemented our analyses with an independent, pedigree-based recombination map. Fine-scale recombination rates were extracted from the linkage map of Rastas *et al*. (2016) for which the coordinates were lifted on the latest three spined stickleback reference genome (Sylvestre *et al*., 2023). To assess concordance between linkage map recombination rate estimates and Pyrho-inferred recombination rates, we computed chromosome-wise Pearson correlation coefficients between recombination rate in cM/Mb (from the linkage map) and mean population-scaled recombination rate (ρ) using the *cor.test* function in R.

### Identification of Highly Differentiated Genomic Regions

We identified regions of elevated genetic differentiation between marine and fluvial populations by calculating per-SNP Weir and Cockerham’s F_ST_ using the *--weir-fst-pop* function in VCFtools. F_ST_ values were aggregated into non-overlapping 100 kb windows, a size selected to balance SNP density and local resolution, given the rapid decay of LD in the three-spined stickleback genome.

To control the confounding influence of recombination rate variation, we assigned each window to one of five recombination bins defined by the 0–20th, 20–40th, 40–60th, 60-80^th^ and 80–100th percentiles of the genome-wide recombination distribution. As discussed previously, the recombination bins were defined with the linkage map recombination rate instead of the Pyrho-inferred recombination rate to avoid circularity. Within each bin, we defined outlier windows as those exceeding the 99th percentile of F_ST_ values, allowing for recombination-informed detection of elevated differentiation without applying a uniform genome-wide threshold.

To further reduce SNP-level noise, we assessed the significance of outlier SNP enrichment per window using a binomial framework. For each 100 kb window, we calculated the binomial probability of observing a given number of outlier SNPs or more, considering the total number of SNPs in the window and the genome-wide proportion of outliers. While non-independence among SNPs precludes formal p-value calculation, this enrichment test highlights windows containing disproportionate numbers of highly differentiated variants. Windows with a non-exact p-value < 0.05 were retained for downstream analyses. Adjacent significant windows were subsequently merged into single continuous regions.

Because variation in F_ST_ can also reflect underlying heterogeneity in nucleotide diversity, we modeled windowed F_ST_ as a function of recombination rate and diversity. Nucleotide diversity (π) was estimated separately in marine and fluvial populations using VCFtools (*--window-pi*) in 100 kb non-overlapping windows. A multiple linear regression by population was then used to test the effect of the linkage map recombination rates and π on F_ST_.

In addition, we assessed whether local mismatches between LD-based recombination estimates and pedigree-based rates coincide with highly differentiated regions. For this, we computed local correlations between log-transformed Pyrho recombination rates (ρ) and linkage map estimates (cM/Mb) using 11-window sliding windows. Each window was assigned to one of five bins (also divided by percentiles of the distribution) based on the absolute correlation value, and we tested for enrichment of highly differentiated regions in each bin using binomial tests against the genome-wide proportion of enriched windows.

Finally, we evaluated the genome-wide association between genetic differentiation and recombination divergence using a Spearman correlation between windowed F_ST_ and PRDI, calculated in non-overlapping 5 Mb windows. Correlation significance was assessed with the *cor.test* function in R.

### Structural Variant Catalogue and Genotyping

We analyzed structural variation using an updated version of the SV catalogue established in Delaive *et al*. (2025), which combines long-read and short-read sequencing data. In the original study, long reads from Oxford Nanopore sequencing were aligned to the stickleback reference genome using Winnowmap v2.03. SVs were identified using three long-read callers (Sniffles2, SVIM, NanoVar) and three short-read callers (Delly, Manta, Smoove). Variants supported by at least two callers within each data type were retained, and long- and short-read sets were merged into a unified SV catalogue. This combined list of deletions, insertions, and inversions was genotyped across all individuals using VG Giraffe, a graph-based genotyping tool designed to account for complex structural polymorphisms. For the present study, we retained the full set of insertions and deletions as defined in the original catalogue.

Inversions were re-evaluated to improve reliability. Initial Sniffles-based inversion calls were manually curated using a protocol developed by Fantine Benoit (in prep.). For two reference individuals (BERG_23 and CHAT_02), SVs were inspected in IGV to assess read support, breakpoint clarity, and overlap with other SVs. Inversions lacking sufficient support, showing ambiguous breakpoints, or overlapping other rearrangements were excluded. Of the 230 inversion candidates examined, 43% were confirmed to be true inversions manually. We further collapsed nested and overlapping variants by retaining only the longest non-overlapping representative variant. This curated inversion set, along with the previously validated insertions and deletions, was re-genotyped across all individuals using an updated genome graph genotyping pipeline (https://github.com/LaurieLecomte/genotype_SVs_SRLR_update.git).

### Analysis of Structural Variants

To identify SVs with elevated differentiation between habitats, we calculated F_ST_ by variant between marine and fluvial populations using VCFtools’ *–weir-fst-pop* function. Outliers were defined as SVs exceeding the 99^th^ percentile of the F_ST_ distribution.

To test whether outlier SVs were preferentially located within highly differentiated genomic regions, we performed a Fisher’s exact test based on their overlap with windows enriched for F_ST_ outlier SNPs. We then constructed a 2×2 contingency table contrasting the number of outlier and background SVs that overlapped at least one F_ST_-enriched window versus those that did not. The Fisher’s exact test was implemented in R using the *fisher.test()* function with a two-sided alternative hypothesis.

We evaluated the impact of SVs on local recombination by modeling patterns of LD around SVs. Pairwise LD (r²) was estimated using the *--geno-r2* function in VCFtools, within 35,000 bp windows, separately for each chromosome and population. For each SV, we extracted a single SNP pair flanking the SV within ±5 kb and calculated three explanatory variables: the inverse of the frequency-weighted physical distance between SNPs, the frequency of the SV in the population, and the proportion of the interval between flanking SNPs spanned by the SV. Weighted distance was defined as the product of the SNP pair distance (including SV length) and SV heterozygosity, the latter estimated from allele frequencies using VCFtools *--freq*. This formulation accounts for the fact that the observed LD reflects the relative abundance of the SV alleles: for instance, if a deletion is frequent, the flanking SNPs will more often be physically closer in the haplotypes carrying it, compared to when the non-deletion allele is more common. We fit linear mixed-effects models separately for the freshwater and saltwater datasets adding the SV type as another explanatory variable and the name of the chromosome as random effect. Both frequency-weighted distance and the flanking SNP interval were scaled prior to modeling. For each population, we modeled r² as a function of scaled distance, proportion covered, and SV type, with chromosome included as a random effect to account for non-independence of observations within chromosomes. This approach allowed us to leverage the full dataset while controlling for chromosome-level structure. Models were fit in R using the lmer function from the lme4 package and estimated marginal means for SV types were compared using the emmeans package.

### Comparison of Repeated and Unique Enriched Windows

To test whether regions previously identified as repeated targets of local adaptation in global stickleback populations (Roberts Kingman *et al*., 2021) overlapped with outlier windows in our dataset, we first lifted over the coordinates of repeated regions to our reference genome using the snplift tool (https://github.com/enormandeau/snplift). Coordinates were manually curated to retain high-confidence mappings.

We then tested for enrichment of overlap between repeated regions and enriched for outlier windows identified previously. Overlap was defined as any intersection between a 100 kb window and a repeated region. To determine whether the observed overlap was greater than expected by chance, we performed a chi-squared test comparing the number of overlapping and non-overlapping windows for both outlier-enriched and non-enriched sets. To account for genome structure and recombination heterogeneity, we also performed a permutation test. At each iteration (n = 500), the set of outlier windows was randomly repositioned across the genome while maintaining chromosome identity and window size. For each permutation, we recalculated the chi-squared statistic and generated a null distribution. An empirical p-value was then computed as the proportion of permuted chi-squared statistics greater than or equal to the observed value.

To further characterize if repeated adaptive regions have a tendency to accumulate in low recombining regions, we compared them to the genomic background for the log-transformed mean recombination rate from pyrho by performing an ANOVA.

## Results

### Population-Specific Recombination Maps

We reconstructed fine-scale recombination maps for marine and fluvial populations of three spined sticklebacks from the St. Lawrence Estuary using a LD-based approach.

To contextualize recombination patterns, we first inferred historical changes in Ne using SMC++. Both populations showed signatures of demographic expansion following an ancestral bottleneck. In the fluvial population, contemporary Ne estimates ranged from 67,192 to 6,513,355, with a median of 1,572,715. In contrast, the marine population showed a narrower contemporary Ne distribution, ranging from 37,031 to 135,695, with a median of 105,445. These estimates reflect relative changes in population size over time.

Genome-wide recombination rates varied substantially across chromosomes and along their lengths in both populations (Figure 2). In the fluvial (freshwater) population, rates ranged from 4.9 × 10⁻¹³ per base per generation on chromosome XI to a maximum of 1 × 10⁻⁵ on chromosome XII. In the marine population, values spanned from 1.2 × 10⁻²³ on chromosome XVI to 1 × 10⁻⁶ on chromosome V. To evaluate the reliability of these LD-based estimates, we compared them with an independent pedigree-based linkage map (Rastas *et al*., 2016). Genome-wide recombination rates from both approaches were strongly correlated (Pearson’s r = 0.664), with most chromosomes showing even higher concordance (r = 0.72–0.77). Notably, correlations were weaker (r ≈ 0.61) on chromosomes V, IX, XI, XVI, and XXI, which also harbored the most extreme recombination values in our populations.

**FIGURE 2.**
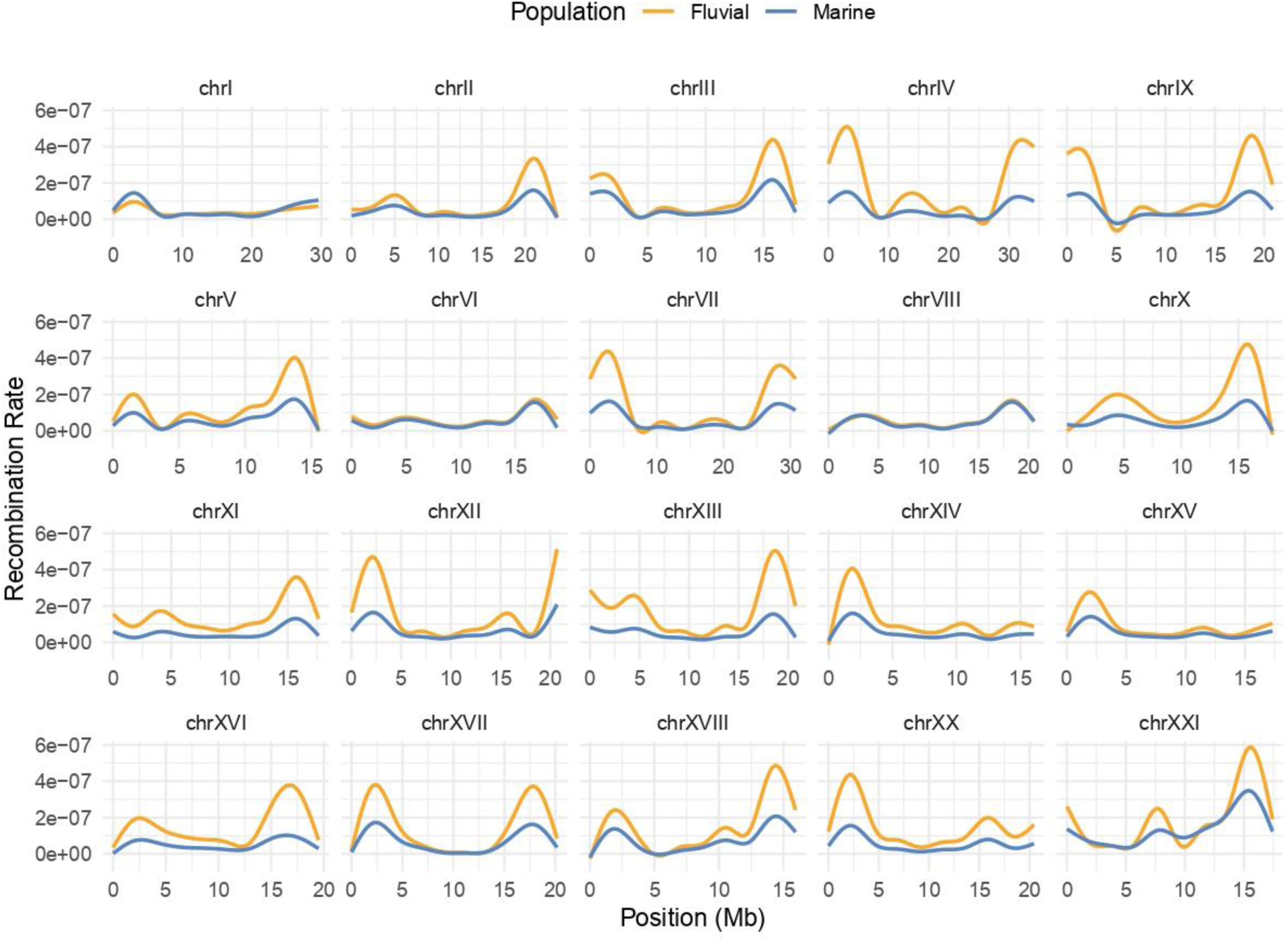
Genome-wide recombination rate variation in freshwater and marine three-spined stickleback. Recombination rates varied substantially across chromosomes and along their lengths in both populations. Each panel shows one chromosome, with the x-axis representing position in megabases (Mb) and the y-axis showing the per-base, per-generation recombination rate. Separate lines indicate each population.

### Divergence in Recombination Landscapes Between Populations

We assessed whether recombination landscapes differed between freshwater and marine populations by comparing recombination rate similarity within and between replicate maps. Pairwise Spearman correlations revealed higher consistency within populations (ρ = 0.91 in freshwater; ρ = 0.90 in marine) than between them (ρ = 0.80), indicating that recombination landscapes were more similar among replicates from the same population than across populations (Figure 3.A).

**FIGURE 3.**
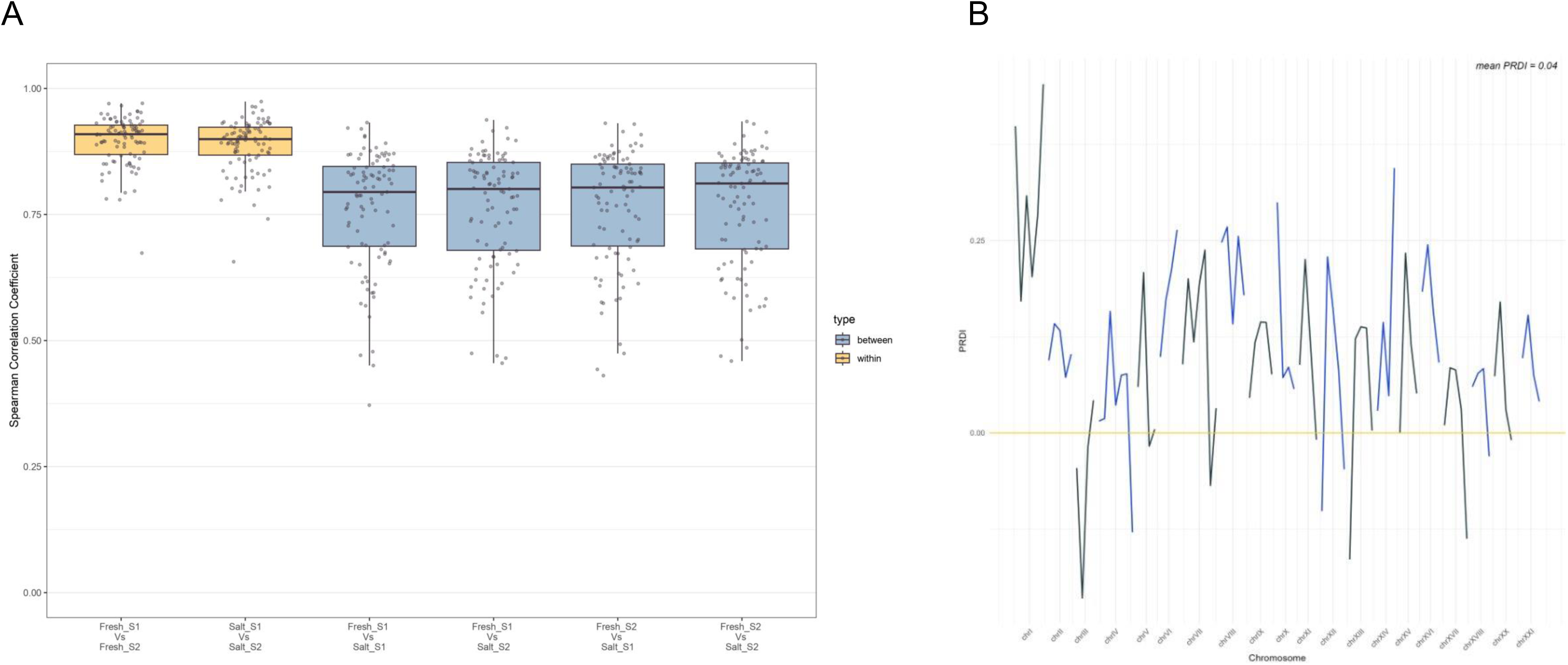
Differences in recombination landscape similarity between freshwater and marine three-spined stickleback. (A) Boxplot representing pairwise Spearman correlations of recombination rates within and between replicate maps for fluvial (ρ = 0.91) and marine (ρ = 0.90) populations, and between populations (ρ = 0.80). Correlations were higher within populations than between them, indicating greater similarity among replicates from the same population. S1 and S2 indicate the two subsets analysed per population. (B) Variation in PRDI along the genome. Positive values indicate genomic regions with greater divergence in recombination landscapes between populations, whereas negative values denote conserved regions.

To determine whether this reduction in between-population correlation could be explained by different demographic history alone, we performed neutral coalescent simulations under a model assuming identical recombination landscapes in both populations. In simulated data, within-population correlations were slightly higher (ρ = 0.95 in freshwater; ρ = 0.93 in marine), and between-population correlation also increased (ρ = 0.88). The difference between empirical and simulated values was summarized using the Population Recombination Divergence Index (PRDI), yielding a genome-wide value of 0.04.

PRDI values varied along the genome, indicating spatial variation in recombination landscape divergence. Positive PRDI values indicate a difference in the recombination landscapes between populations while negative values point to more conserved regions. The highest value was observed on chromosome I (PRDI = 0.45), while the lowest occurred on chromosome III (PRDI = –0.21) (Figure 3.B).

To further quantify localized divergence, we computed pairwise recombination rate dissimilarity (Δ*r*) between freshwater and marine populations in non-overlapping 100 kb windows. This analysis identified no windows with elevated between-population divergence relative to within-population variation. When lowering the threshold to two standard deviations over the mean, we identified 15 outlier windows. Of these, 12 were located on chromosome XVII.

We next examined fine-scale recombination features by identifying hotspots and coldspots of recombination in each population. Using a deviation-based thresholding approach with a 40 kb background, we detected 70 recombination hotspots in the marine population and 120 in the fluvial population. Among these, 25 hotspots were shared between the two populations (13% overlap, representing 36% of marine and 21% of freshwater coldspots). Coldspots were more consistently shared: 129 were identified in the marine population and 134 in freshwater, with 76 shared (29% overlap, representing 59% of marine and 57% of freshwater coldspots).

### Recombination Landscape Influence Patterns of Genomic Differentiation

To evaluate how recombination rate variation contributes to genetic differentiation, we identified 553 windows enriched for highly differentiated SNPs using a recombination-informed framework. These enriched windows were distributed evenly across five recombination bins and spanned 0.02% of the genome.

When comparing recombination estimates, genome-wide correlations between Pyrho-inferred and linkage map recombination rates were generally high but dropped locally, sometimes in regions overlapping enriched windows. However, binomial enrichment tests across five correlation bins revealed no significant association between the magnitude of local correlation and the likelihood of detecting an F_ST_ outlier (all p > 0.19), indicating that local mismatches between recombination maps are not systematically predictive of differentiation.

While investigating the broad relationship between genomic differentiation and recombination divergence, we observed a positive, though marginally non-significant association between PRDI and F_ST_ values across 5 Mb windows (Spearman ρ = 0.274, p = 0.0654; Figure 4), suggesting a tendency for highly differentiated regions to exhibit greater recombination divergence.

**FIGURE 4.**
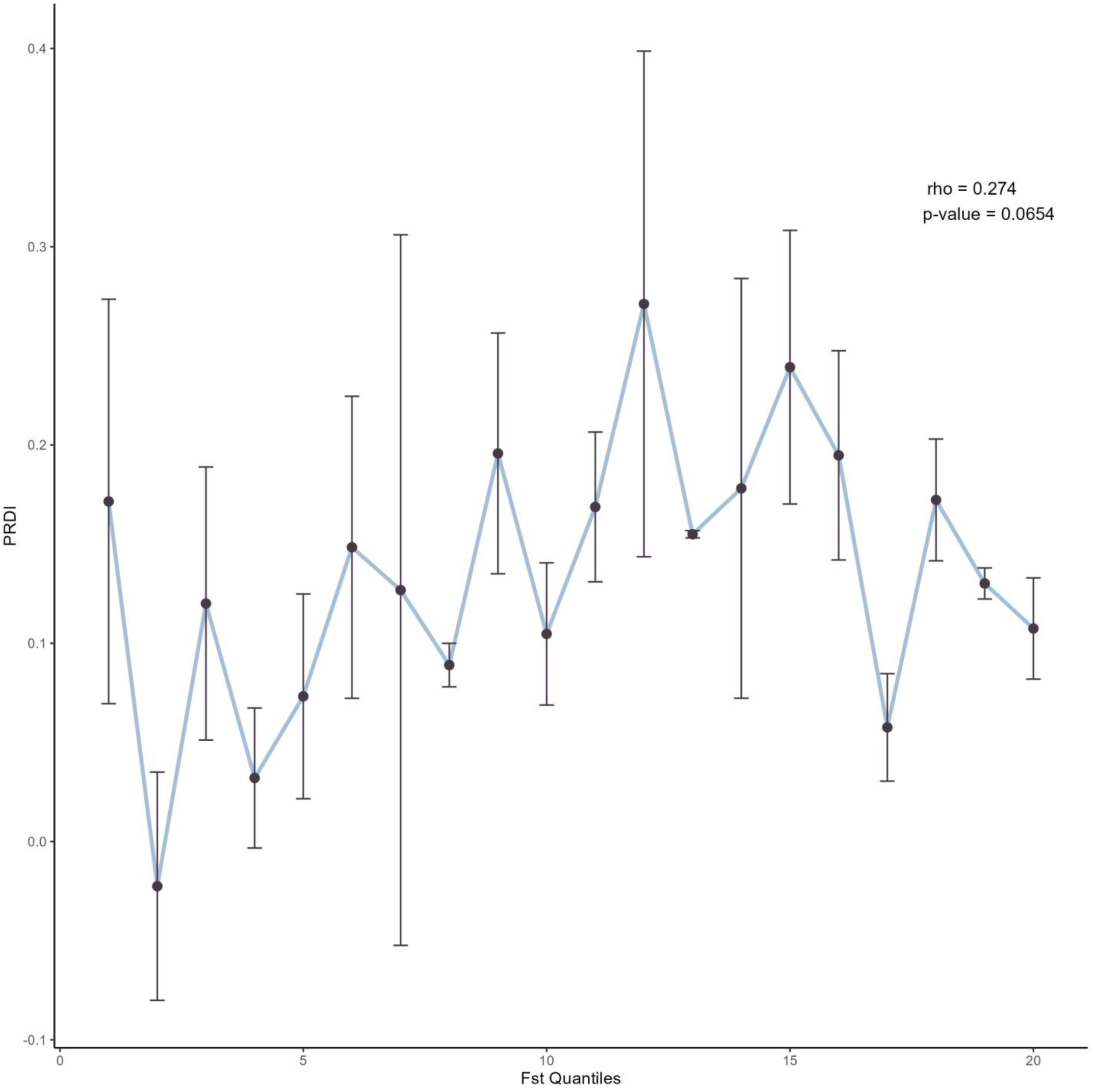
Relationship between genomic differentiation and recombination landscape divergence in three-spined stickleback. Scatterplot showing the association between PRDI and F_ST_ quantiles, where F_ST_ was calculated on 100 kb windows. Each point represents the mean PRDI within a given F_ST_ quantile, with error bars indicating 95% confidence intervals. Across 5 Mb genomic windows, PRDI and F_ST_ were positively, though marginally non-significantly, correlated (Spearman ρ = 0.274, p = 0.0654), suggesting a tendency for highly differentiated regions to exhibit greater recombination divergence.

Because elevated F_ST_ may result from both directional selection and confounding factors such as linked or background selection or ancestral diversity differences, we modeled log-transformed F_ST_ as a function of recombination rate and within-population nucleotide diversity (π), separately for the freshwater and saltwater populations. Both models explained a similar proportion of F_ST_ variance (adjusted R² = 0.134 for freshwater; 0.131 for saltwater, p < 2.2 × 10⁻¹⁶ in both cases), and in each case revealed a significant interaction between π and recombination rate (interaction estimates = −7.98 and −7.57, p < 1 × 10⁻⁵). These interactions indicate that the relationship between nucleotide diversity and genetic differentiation depends on the recombination landscape: specifically, the association between π and F_ST_ is stronger in regions of low recombination.

These results suggest that part of the observed differentiation may arise from variation in nucleotide diversity, particularly in low recombination regions where linked selection is expected to be most influential.

### Structural Variants impact the recombination landscape

We identified a total of 37,935 SVs distributed across the genome, comprising 19,790 deletions, 18,103 insertions, and 42 inversions. Deletions had a mean length of 216 bp (range: 142–24,911 bp), while insertions were slightly larger on average (mean = 292 bp, range: 79–15,792 bp). Inversions were less frequent but considerably longer, with a mean size of 3,117 bp and lengths ranging from 47 to 28,118 bp.

To identify habitat-associated SVs, we calculated F_ST_ between marine and fluvial populations and defined outliers as those exceeding the 99th percentile of the F_ST_ distribution. This resulted in 386 outlier SVs, comprising both deletions and insertions. No inversions met the outlier threshold, likely due to their limited sample representation.

To assess whether outlier SVs were disproportionately located within highly differentiated genomic regions, we tested their overlap with windows enriched for F_ST_ outlier SNPs. A large majority of outlier SVs (78.2%) overlapped with at least one F_ST_ outlier window, compared to 21.8% of background (non-outlier) SVs. This enrichment was highly significant (Fisher’s exact test, *p* = 2.35 × 10⁻¹²¹) with an odds ratio of 12.90, indicating that outlier SVs were nearly 13 times more likely to occur within regions of elevated genetic differentiation based on SNPs.

We modeled LD (r²) around SVs using linear mixed-effects models that included SV type, distance-weighted SNP separation, and interval between flanking SNPs as fixed effects, with chromosome as a random effect to account for chromosome-level variation. In the freshwater dataset, the interval between flanking SNPs was significantly associated with r² (β = 0.0085, *t* = 12.97), while the effect of frequency-weighted distance was negligible (β = 4.6 × 10⁻ ⁵, *t* = 0.07). Inversions were associated with significantly higher LD compared to both deletions (Δr² = 0.0519, *p* = 0.0317) and insertions (Δr² = 0.0532, *p* = 0.0266), as indicated by pairwise comparisons of estimated marginal means. LD around deletions (mean r² = 0.0518) and insertions (mean r² = 0.0505) did not differ significantly (*p* = 0.5847). In the saltwater dataset, the interval between flanking SNPs also had a strong positive effect on r² (β = 0.0094, *t* = 15.42), while distance had no significant effect (β = –0.00020, *t* = –0.33). Although LD around inversions appeared elevated (mean r² = 0.0626) relative to deletions (0.0398) and insertions (0.0380), these differences were not statistically significant (all *p* > 0.30).

### Overlaps with Globally Repeated Adaptive Regions

To evaluate whether locally enriched genomic regions in our population also coincide with previously identified genomic regions exhibiting repeated signatures of adaptation across the species range, we compared our 100 kb windows enriched for FST outliers with the 92 outlier regions identified by Roberts Kingman *et al*. (2021; hereafter “repeated region”). Among the 1,037 enriched windows identified in this study, 71 (6.85%) overlapped at least one repeated region (Figure 5). Conversely, 54 of the 92 previously identified repeated regions (58.6%) contained at least one enriched window. This represents a more than twofold enrichment compared to background genomic windows, 3.03% of which overlapped with repeat regions (χ² = 30.67, p < 0.001). A permutation test confirmed that this enrichment is unlikely to occur by chance, with none of the 500 permuted χ² values exceeding the observed value (mean = 1.53, SD = 2.05, empirical p < 0.002).

**FIGURE 5.**
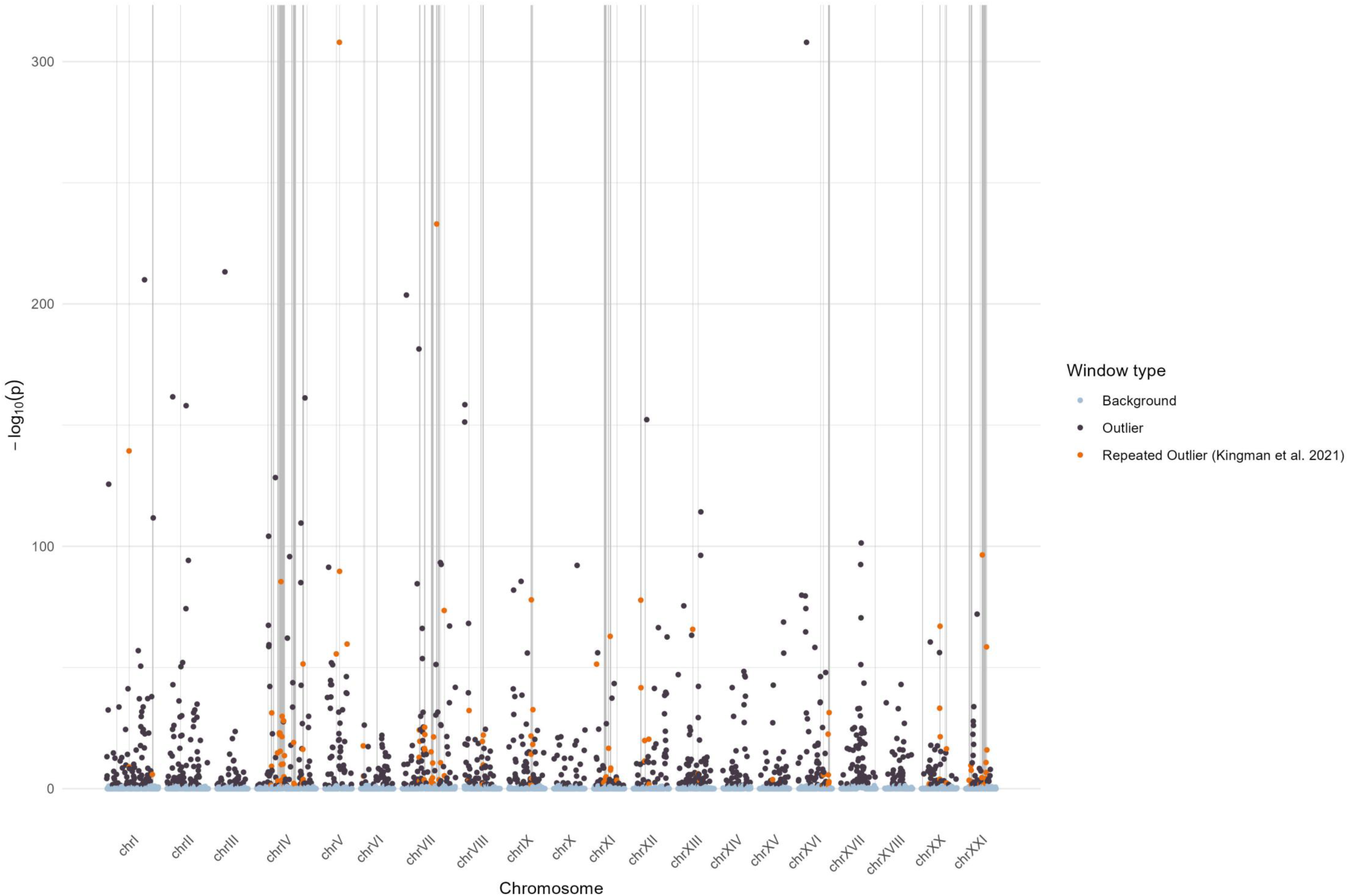
Overlap between enriched regions in our dataset and globally repeated targets of adaptation in three-spined stickleback. Each point represents a 100 kb genomic window, with light grey regions corresponding to the 92 adaptive regions identified by Kingman et al. (2021). Enriched windows are shown in black, and those located within grey regions, termed “repeated outliers”, are shown in orange. The y-axis represents the –log_10_-transformed non-exact *p*-value from a binomial test assessing enrichment of outliers in each window. Among 1,037 enriched windows, 71 (6.85%) overlapped at least one repeated region, and 54 of the 92 repeated regions (58.6%) contained at least one enriched window.

Repeated regions were broadly distributed across the genome and recombination landscape. Nevertheless, a significant decrease in the mean recombination rate estimated with Pyrho was observed in enriched windows overlapping with repeated regions compared to enriched windows not overlapping repeated regions (mean r^2^_non-repeated_ = 7.8 × 10⁻ ^8^ vs mean r^2^_repeated_ = 3.9 × 10⁻ ^8^, p < 2 x 10^-16^). SVs covered 2.07% of the repeated regions, with 1,681 SVs identified across these loci, including 920 insertions, 759 deletions, and two inversions on chromosome I and VII. Of those overlapping SVs, 56 of them were outlier SVs. These findings suggest that globally reused regions of adaptation are concentrated in lower recombination regions and harbor substantial structural polymorphism.

## Discussion

Understanding how recombination landscapes and structural variation interact to shape the genomic architecture of local adaptation under gene flow is central to explaining how adaptive divergence persists in nature. In this study, we compared two geographically close but ecologically distinct populations of three-spined sticklebacks in the St. Lawrence Estuary to test how recombination variation influences the retention of locally adaptive alleles. We found that recombination landscapes varied markedly between populations, revealing population-specific patterns rather than a shared genomic template. Contrary to expectations that local adaptation should be enriched in low-recombination regions, adaptive divergence occurred across regions of varying recombination rates. SVs, particularly insertions, deletions and inversions, emerged as key modifiers of these recombination landscapes, influencing where and how adaptive regions are maintained. In the presence of persistent gene flow, these processes resulted in a more concentrated genomic architecture of local adaptation, with repeatedly implicated regions from Roberts Kingman *et al*. (2021) contributing disproportionately to adaptive divergence.

Overall, our findings highlight the complex interplay between recombination, structural variation, and selection in shaping adaptive divergence between connected populations, and underscore the value of integrating recombination mapping with SV analyses to understand the genomic basis of local adaptation in natural populations.

### Recombination landscapes diverge between populations

We found that recombination landscapes differ meaningfully between marine and fluvial populations of three-spined stickleback. In particular, the fluvial population exhibited higher overall recombination rates, potentially reflecting a larger Ne in the fluvial population. While earlier inferences based on fastsimcoal pointed to the opposite trend (Delaive *et al*., 2025), the discrepancy likely stems from methodological differences. Unlike fastsimcoal, which relies on predefined demographic models, SMC++ reconstructs Ne trajectories through time without assuming specific scenarios. Under this framework, the higher Ne in freshwater likely reflects a stronger or more recent post-bottleneck expansion, which aligns with expectations for recently colonized freshwater habitats. However, it should be noted that SMC++ assumes no migration and can be sensitive to population structure, so the absolute Ne estimates (particularly the relatively low Ne inferred for the marine population) should be interpreted cautiously and may reflect local sampling rather than the full marine metapopulation.

Despite this difference in magnitude, the overall structure of the recombination landscape remains largely conserved, with genome-wide recombination rates showing strong correlation between populations (r ≈ 0.8). Still, subtle but significant divergence was detected using the PRDI metric. When estimating Ne through time, SMC++ assumes no gene flow in the population. While this assumption does not reflect the low but real gene flow documented in the system, we simulated recombination landscapes under a model with no migration between populations. Notably, Talbi *et al*. (2025) showed that increasing migration in simulations artificially inflates PRDI values. Therefore, the PRDI values we report likely represent a conservative estimate of recombination landscape divergence between these partially connected populations.

Interestingly, PRDI appeared more sensitive to recombination landscape divergence than the Δr metric. This likely reflects the nature of the metrics themselves: while Δr quantifies absolute differences in recombination rates, PRDI tracks relative shifts. The weaker signal in Δr suggests that although landscapes are reshaped between populations, these changes may not be large in magnitude.

The most dynamic aspect of the recombination landscape appeared at fine scales. Only 39% of recombination hotspots were shared between populations, suggesting rapid hotspot turnover—a pattern previously reported in sticklebacks (Shanfelter, Archambeault and White, 2019) and consistent with recent findings in three salmonid species (Raynaud *et al*., 2025). This high turnover is thought to stem from the inherently unstable nature of hotspot determinants, such as sequence motifs or chromatin accessibility (Jeffreys and Neumann, 2009). Our findings support this interpretation and are consistent with the “hotspot paradox,” whereby recombination depletes its own targets over time, shifting activity to nearby regions (Úbeda, Fyon and Bürger, 2023).

### Interaction between genomic differentiation and recombination

Candidate outlier regions identified through our recombination-informed enrichment framework encompassed ∼0.02% of the genome. These regions were distributed relatively evenly across recombination bins, indicating that elevated differentiation was not strongly biased toward a particular recombination environment. While theoretical regions suggest that low recombination regions should facilitate the buildup of differentiation by reducing the homogenizing effect of gene flow and preserving locally beneficial haplotypes (Tigano and Friesen, 2016; Marques, Meier and Seehausen, 2019), differentiation can still occur without such facilitation, and these observations show that any such effect may be weak or subtle in natural populations.

The lack of such a pattern in our data likely reflects the polygenic and functionally diverse nature of freshwater adaptation in three-spined sticklebacks. This process involves a broad suite of traits, including morphology, behavior, respiration, and predator defense, highlighting its functional diversity (Schluter, 1993; Taugbøl *et al*., 2020; Aguirre *et al*., 2022). As a result, selection can act across numerous genomic regions, potentially under different selection intensities and histories. Strongly selected alleles that arose early in the adaptive process may now be found in regions with high recombination rate, where the benefits of recombination (such as reducing linkage with deleterious mutations) outweigh the advantages of maintaining linkage with co-adapted alleles (Otto and Feldman, 1997; Charlesworth, 2012). In contrast, more recently selected alleles or those involved in complex epistatic interactions may be preferentially retained in low recombination regions, where linkage preserves beneficial allele combinations (Parée *et al*., 2025). This duality could explain the lack of a simple relationship between recombination and outlier enrichment. Alternatively, the influence of recombination on outlier enrichment may be relatively weak compared to the strength of selection. If adaptation requires changes at specific loci, these targets can still accumulate adaptive mutations even when they fall in regions of higher recombination, leading to detectable signals of differentiation irrespective of local recombination context.

These findings also align with the concerns raised by Booker, Yeaman and Whitlock (2020), who warned that ignoring recombination rate heterogeneity in genome scans could lead to biased inference of selection. Specifically, F_ST_ distributions tend to have long tails in low recombination regions, increasing the propensity to fall outside of any threshold based on genome-wide values, with the opposite pattern in high recombination regions due to their tighter distributions. Our results confirm this pattern, underscoring the importance of accounting for recombination when interpreting genome-wide scans of differentiation

Interestingly, although the average recombination rate was not predictive of differentiation, we did detect a relationship between PRDI (a measure of recombination rate divergence) and F_ST_. In our dataset, higher FST values tended to be associated with greater differences in nucleotide diversity between populations. This likely reflects the impact of linked selection, whereby regions of low diversity often exhibit elevated LD, leading to downward-biased recombination estimates, whereas regions of high diversity produce less biased but noisier estimates. This circularity suggests that observed correlations between PRDI and F_ST_ could partly reflect underlying differences in genetic diversity, especially in low recombination regions where both linked selection and ancient coalescent events can exacerbate differentiation (Noor and Bennett, 2009; Cruickshank and Hahn, 2014). Comparable links between recombination, diversity, and differentiation have been observed in Eurasian blackcap where population-specific reduction of recombination biased their detection of potentially adaptive variants (Ishigohoka *et al*., 2024).

Altogether, these observations support a more nuanced view of how recombination influences the genomic landscape of divergence. Rather than a uniform bias toward differentiation in low recombination regions, our results suggest that the strength and timing of selection, the haplotype structure of a genomic region, the polygenic nature of adaptation, and variation in genetic diversity all contribute to determine where outliers emerge in the genome.

### Structural variants modulate the recombination landscape

The patterns observed in our study support a role for SVs in shaping the recombination landscape. We found that SNP pairs for which a larger proportion of their distance was spanned by an SV exhibited elevated LD, consistent with recombination suppression across multiple SV categories. Similar patterns have been reported elsewhere: Morgan *et al*. (2017) found that recombination coldspots in mice often coincided with CNVs, while Huang *et al*. (2025) showed that transposable elements generating structural variation also suppressed recombination in their vicinity in *Drosophila melanogaster*. Together, these findings reinforce the idea that SVs may interfere with chromosome pairing or recombination initiation, which may be driven by both structural constraints and epigenetic mechanisms (Casale *et al*., 2024; Johnston, 2024).

The strongest effect of SV on recombination was detected for inversions, which substantially increased LD and thus likely reduced effective recombination rates. Interestingly, this relationship was statistically significant only in the fluvial population, although a similar tendency was present in the marine population. Several explanations are possible. On the technical side, the larger sample size for the marine group (n = 270 vs. 89) may have increased the precision of LD estimates but also made them more sensitive to subtle population substructure. Such substructure can artificially inflate LD by creating correlations in allele frequencies among groups, even in the absence of reduced recombination (Li and Nei, 1974; Ohta, 1982). This in turn reduces the accuracy of recombination rate inference and makes it more difficult to disentangle the effect of SVs from background demographic structure. Biologically, factors such as inversion age, size, or associated haplotype composition can influence the extent of recombination suppression (Stevison, Hoehn and Noor, 2011; Charlesworth, 2023). However, given that inversion heterozygosity was similar in both populations, these factors alone may not account for the observed difference.

Together, these results suggest that SVs are important contributors to variation in recombination rates and may shape patterns of genetic variation in both connected and partially isolated populations. Inversions showed the strongest and most consistent recombination-suppressing effects, but other SV types should not be overlooked. Experimental work has shown that insertions and deletions can alter recombination outcomes at the molecular level; for example, the presence of a gap, deletion, or heterologous insertion in DNA templates markedly reduced the efficiency of recombination compared to intact homology (Brenner, Smigocki and Camerini-Otero, 1985). Additionally, CNV and intrachromosomal rearrangements have been linked to recombination rate variation in birds, where CNVs and rearrangements were enriched in regions of elevated recombination (Völker *et al*., 2010). Although the effects of insertions and deletions in our dataset were more modest than those of inversions, these findings indicate that multiple SV categories, acting through both molecular and structural mechanisms, can influence recombination landscapes. This highlights the importance of considering the full spectrum of SVs, rather than focusing exclusively on inversions, when interpreting the genomic architecture of recombination.

### Concentrated Genomic Architecture of Local Adaptation in the Presence of Gene Flow

The detection of repeat regions shared between our dataset and the globally reused targets of adaptation reported by Roberts Kingman *et al*. (2021), together with previous work in the St. Lawrence Estuary by McCairns and Bernatchez (2008), supports the presence of local adaptation despite ongoing gene flow. McCairns and Bernatchez showed that fluvial and marine sticklebacks in this system form distinct genetic groups and that environmental rather than geographic factors explain a large proportion of their differentiation, highlighting the role of ecological selection in maintaining divergence under migration. Our results extend this view by demonstrating that several of the same genomic regions involved in global freshwater adaptation also contribute to divergence in the St. Lawrence Estuary. This reuse of previously identified globally repeated targets of local adaptation is consistent with the species’ well-documented capacity for rapid adaptation to freshwater (Barrett *et al*., 2011; Terekhanova *et al*., 2014). At the same time, our results highlight a suite of genomic regions unique to our system, which may reflect ecological differences in the St. Lawrence Estuary compared to more typical freshwater–marine contrasts. For instance, unlike many freshwater stickleback populations, our fluvial fish retain extensive armor, likely due to the persistence of predators in the estuary (García-Machado *et al*., 2022) as it is the case for other estuarine stickleback populations from western Canada (Reimchen, 2000). Nonetheless, other traits such as feeding ecology, reproductive behavior, and aspects of morphology do differ between environments (Mccairns and Bernatchez, 2012), as do abiotic variables such as pH, salinity, and temperature. Genes associated with these traits are possibly those contributing to the repeated signals we observed.

Repeated regions were not randomly distributed across the genome. Instead, more than half (54%) of them were concentrated on three chromosomes (IV, VII and XXI), which we designated as major hotspots of local adaptation in this system. This pattern is consistent with previous reports of concentrated architectures in stickleback (Hohenlohe *et al*., 2010; Jones *et al*., 2012; Roberts Kingman *et al*., 2021). These regions may represent clusters of tightly linked adaptive loci, as suggested by their reduce recombination rates compared to non-repeated outlier windows, or they may house highly pleiotropic genes under strong selection. Such a combination of clustered and distributed patterns is in line with theoretical expectations under migration-selection balance, where selection maintains large-effect loci in low recombination regions while smaller-effect loci are distributed more broadly across the genome (Yeaman and Whitlock, 2011; Yeaman, 2022). Indeed, a more concentrated architecture of local adaptation was found in *Populus trichocapra* populations that were subject to higher gene flow compared to more distant populations (Holliday *et al*., 2016).

Interestingly, none of the 42 inversions we genotyped showed differentiation high enough to qualify as outliers, an unusual result given the prominence of inversions in many recent studies of adaptation with gene flow (Shi *et al*., 2023; Fuentes-Pardo *et al*., 2024; Sabatino *et al*., 2025). This may indicate that inversions in our system are relatively young, as suggested by their lower F_ST_ and minimal variation in nucleotide diversity (Akopyan *et al*., 2025). Still, their potential role cannot be ruled out as two inversions, on chromosomes I and VII, overlapped repeated adaptive regions. The former has been previously linked to freshwater adaptation in sticklebacks (Rodríguez-Fuentes *et al*., 2025), while the latter, associated with defense trait QTL (Peichel and Marques, 2017), is a novel candidate in our system. Notably, we did not detect the known inversions detected in other three-spined stickleback systems on chromosomes XI and XXI (Jones *et al*., 2012; Roesti *et al*., 2015).

Overall, recombination landscapes and SVs, including inversions, insertions, and deletions, influence local patterns of recombination and differentiation. Clusters of repeated adaptive regions coincide with some SVs and low-recombination areas, highlighting how these genomic features contribute to the organization of adaptive variation in connected populations.

## Data availability

Raw sequence reads come from Delaive *et al*. (2025) study and are deposited in the SRA (BioProject PRJNA1327819).

## Acknowledgment

I would like to thank L. Bernatchez, my former PhD supervisor, who helped me design this project, secure the grant, and guided me through the early stages of my scientific career, but who has sadly passed away.

We also would like to thank C. Otis and B. Boyle for the long-read sequencing as well as B. Bougas for her coordination of the project. We also thank X. Dallaire for his contribution to SV analysis, L. Lecomte for the bioinformatic support and A. Xuereb for her help with the manuscript. This research was funded by the Natural Sciences and Engineering Research Council of Canada under the RGPIN-2020-04282 subvention. The project is part of the Ressources Aquatiques Québec (RAQ) research program.

## Author Contributions section

N. D. revised the manuscript and assumed the supervisory role following the passing of Louis Bernatchez. S. D. prepared the genomic libraries for whole-genome sequencing, performed the analysis and wrote the draft of the manuscript. S. Y. provided valuable suggestions and assisted with coding and result interpretation.

